# Kinesin-1 and dynein use distinct mechanisms to bypass obstacles

**DOI:** 10.1101/624056

**Authors:** Luke S. Ferro, Sinan Can, Meghan A. Turner, Mohamed M. Elshenawy, Ahmet Yildiz

## Abstract

Kinesin-1 and cytoplasmic dynein are microtubule (MT) motors that transport intracellular cargos. It remains unclear how these motors move along MTs densely coated with obstacles of various sizes in the cytoplasm. Here, we tested the ability of single and multiple motors to bypass synthetic obstacles on MTs *in vitro.* Contrary to previous reports, we found that mammalian dynein is highly capable of bypassing obstacles. Unlike dynein, single kinesin motors stall in the presence of obstacles, consistent with their inability to take sideways steps to neighboring protofilaments. Kinesins overcome this limitation when working in teams, bypassing obstacles as effectively as multiple dyneins. Cargos driven by multiple kinesin or dyneins are also capable of rotating around the MT to bypass large obstacles. These results suggest that multiplicity of motors is required not only for transporting cargos over long distances and generating higher forces, but also for maneuvering of the cargos on obstacle-coated MT surfaces.

## Introduction

Kinesin and dynein move towards the plus- and minus-ends of MTs, respectively, and play major roles in intracellular cargo transport, cell locomotion and division.^1, 2^ Although these motors have complementary functions on MTs, they have distinct structural and mechanistic features. Kinesin-1 contains a globular motor domain that binds the MT and hydrolyzes ATP. Two identical motor domains are connected by a short neck-linker to a common tail.^2^ *In vitro* studies have shown that kinesin moves by coordinated stepping of its motor domains, in a manner akin to human walking.^3, 4^ It follows a single protofilament track on the MT and almost exclusively steps forward without frequent sideways or backwards motion.^5, 6^ Unlike kinesin, dynein’s motor domains are large heterohexameric rings of AAA+ ATPase subunits that connect to the MT through a coiled-coil stalk.^7^ Stepping of the dynein motor domains is not tightly coordinated.^8^ Instead, either monomer can take a step while the other serves as a MT tether.^1, 8^ Dynein has a large diffusional component in its stepping behavior, resulting in frequent sideways and backwards steps.^8^ It remains unclear whether the differences in stepping behaviors between these motors influence their cellular functions.

Intracellular transport takes place in a highly crowded and dynamic cytoplasm. The MT network is densely decorated with obstacles such as MT-associated proteins (MAPs), stationary organelles, protein aggregates, MT defects, opposing motor traffic and other cytoskeletal filaments.^9–11^ It is not well understood how motors transport cargos efficiently throughout the cell despite these challenges. Previous *in vitro* studies suggested that motors need to take sideways steps in order to avoid obstacles on their path.^10^ In agreement with this idea, kinesin motility is strongly inhibited by obstacles such as catalytically inactive motors, MAPs or cell extract.^9, 12, 13^ While these motors can occasionally bypass obstacles by detaching and reattaching to a neighboring protofilament, they stall or detach from the MT in most cases.^12, 14^ Kinesin-2, a different kinesin family member with faster MT detachment/reattachment kinetics^15^ and increased side-stepping ability^16^, bypasses obstacles more successfully than kinesin-1. Dynein was expected to be less sensitive to obstacles than kinesin because of its elongated structure and frequent sideways stepping.^8^ Yet, *in vitro* studies on isolated mammalian dynein observed that the motor reverses direction when encountering a MAP obstacle rather than bypassing it.^9, 17^ However, these studies were conducted before it was understood that mammalian dynein alone is autoinhibited and its activation requires assembly with dynactin and a cargo adaptor.^18, 19^ Therefore, it remains unclear whether active dynein motors can bypass obstacles on an MT.

Unlike helicases, unfoldases and DNA/RNA polymerases which usually function as individual motors,^20^ cytoskeletal motors often operate in teams to transport a cargo.^21–23^ Multiple motors carry cargos with increased processivity relative to single motors,^24^ which is essential for long-range transport in cell types such as neurons. Teams of motors also exert higher forces, which may enable transport of large cargos through the dense cellular environment.^25^ It has been proposed that cargos with multiple motors also avoid obstacles more effectively than single motors.^26^ A recent study found that multi-kinesin cargos pause at MT defects rather than detach like single motors.^11^ Ensembles of two kinesin-1 motors linked together with a DNA scaffold have a higher run length than single kinesins, but their run length was also decreased in the presence of neutravidin obstacles on the MT.^15^ Motility of multiple dyneins has not been studied on MTs decorated with obstacles. Therefore, it is not well understood whether cargos driven by multiple motors bypass obstacles more successfully than single motors.

Here, we challenge single- and multi-motor cargos of kinesin and dynein with quantum dot (QD) obstacles on MTs. We find that kinesin and dynein employ different mechanisms to bypass these obstacles. Consistent with their ability to take side-steps, single dynein motors efficiently circumnavigate around QD obstacles. Unlike dynein, single kinesins are strongly inhibited by QDs, yet overcome this limitation when working together as a team. Multiplicity of motors was also critical to bypass large roadblocks. When cargos driven by a team of kinesin or dynein motors face a wall along their path, they swing around the MT without net forward movement and continue moving forward. Together, our results provide insight into how motors transport intracellular cargos along densely-coated MT surfaces.

## Results

### Single dyneins, but not kinesins, avoid obstacles

Previous studies used rigor motors^12^, cell extracts^13^ or MAPs^9^ to study how motors move in the presence of obstacles. Because some of these obstacles have complex binding kinetics to MTs, vary in size, and interact with motors directly, it is difficult to discern how their presence on the MT obstructs motility. We sought a model obstacle that stably attaches to the MT, has a well-defined size and forms no specific interactions with motors. We decorated biotinylated MTs with streptavidin-conjugated QDs (25 nm in diameter).^8^ These QDs have a bright and photostable fluorescent emission, which enable us to measure their linear density along MTs.

We used three motor complexes to study how QD obstacles affect motility: human kinesin-1, yeast cytoplasmic dynein and the mammalian dynein/dynactin/BicD2N (DDB) complex. Motors were labelled with QDs of a different color on their tail region and their motility on surface-immobilized MTs was monitored using multicolor imaging (Fig. 1a). Surface density of QD obstacles was varied, with a maximum decoration of 12 QDs μm^−1^ (Fig. 1b-c). We found that single yeast dynein and DDB motors walked processively even at the highest QD density tested (Fig. 1d-e, Supplementary fig. 1, Supplementary video 1), consistent with dynein’s ability to take side-steps.^8^ We did not see evidence of motor reversals when DDB encountered an QD obstacle, consistent with recent studies of DDB motility on Tau-coated MTs.^27^ However, mobile fraction, velocity and run length of yeast dynein were reduced 30-70% by increasing density of QDs. In comparison to dynein motors, kinesin motility was severely affected by the QDs (Fig. 1d-e). The majority of kinesins became stuck on the MT by addition of QDs (Fig 1d). At low QD density (1-2 μm^-1^), mobile fraction was reduced 90%, while the run length and velocity were reduced by 60% compared to no obstacle condition (p = 0.0003, Fig. 1e, Supplementary video 2). Kinesin motility could not be analyzed at higher QD densities because we did not detect processive runs longer than 250 nm under these conditions. Collectively, these results show that kinesin remains bound to an MT but is unable to move forward when it encounters an obstacle.

**Fig. 1.**
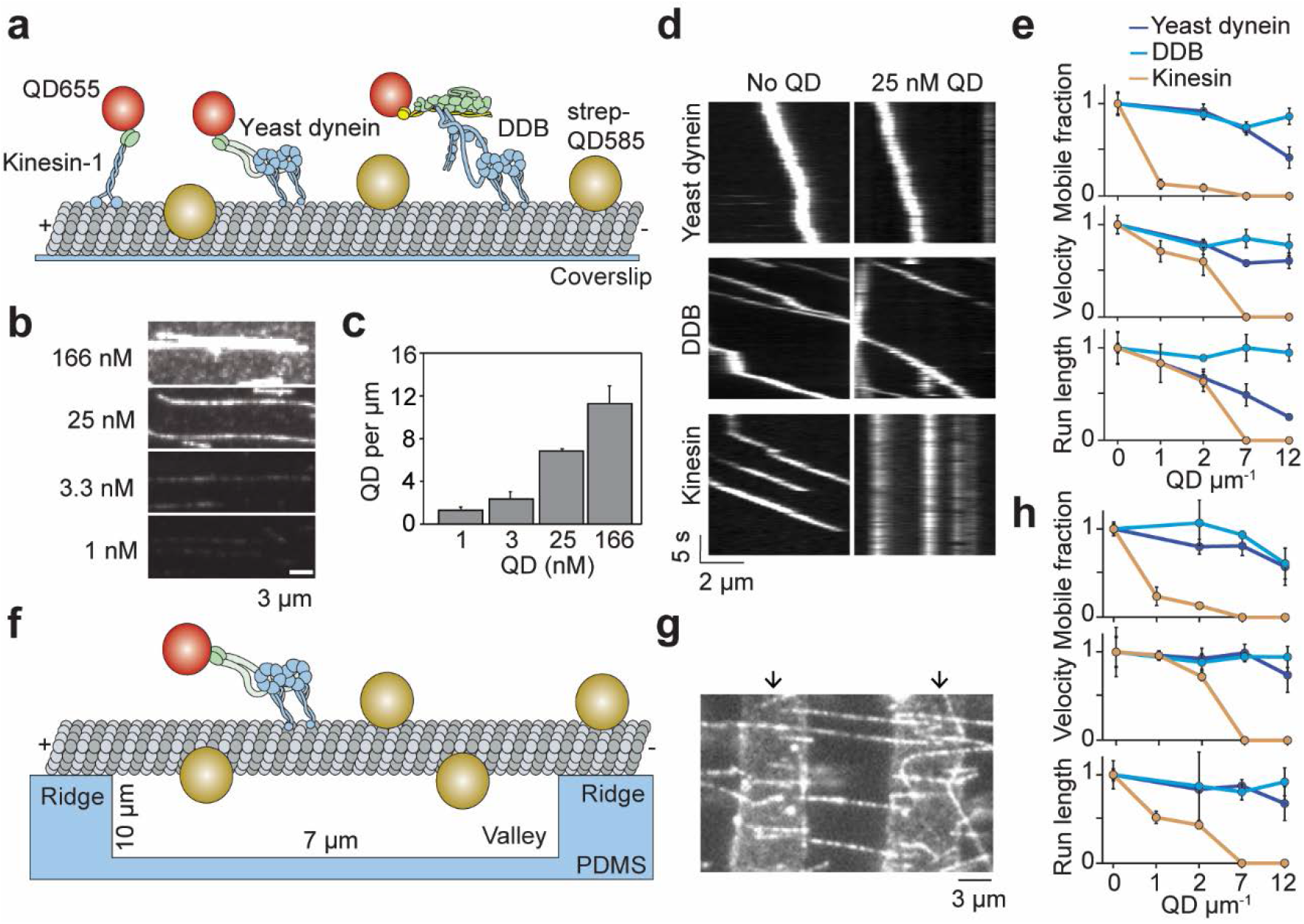
Single dynein, but not kinesin, motors bypass QD obstacles. **a** Schematic of single-molecule motility assays on surface-immobilized MTs decorated with streptavidin-coated QD585 obstacles. Human kinesin-1, yeast dynein and DDB are labeled with QD655 at their tail domain. **b** Example fluorescent images of QD585 obstacles on MTs at different QD concentrations. **c** Linear density of QDs on MTs at different QD concentrations (mean ± SD, from left to right *n* = 97, 98, 104 and 90 MTs from two technical replicates). **d** Kymographs show the motility of QD655-labeled motors on MTs with or without QD obstacles. QD585 signal is not shown. **e** Mobile fraction, velocity and run length for all three motors were normalized to the no QD condition (mean ±SD, three independent experiments). Run length values represent decay constants derived from a single exponential decay fit. From left to right, *n* = 271, 423, 405 for kinesin, 315, 407, 197, 168 for yeast dynein, and 636, 502, 356, 509 for DDB. f Schematic of a single-molecule motility assay on MT bridges coated with QD obstacles (not to scale). g Example image of Cy5-labeled MT bridges in the microfabricated chamber. PDMS ridges (arrows) are visible due to the autofluorescence. h Mobile fraction, velocity and run length of motors along MT bridges were normalized to no QD condition (mean ±SD, three independent experiments). From left to right, *n* = 199, 187, 106 for kinesin, 129, 107, 163, 135 for yeast dynein, and 192, 206, 330, 276 for DDB.

On surface-immobilized MTs, motors cannot access protofilaments facing the coverslip. As a result, surface immobilization may serve as an additional obstacle as the motors attempt to bypass the QDs. MTs in the cell, however, are freely suspended in 3D. This may allow motors to fully explore the MT surface and more successfully bypass the obstacles. To test this possibility, we constructed “MT bridges” by immobilizing MT ends to polydimethylsiloxane (PDMS) ridges on either end of a 10-μm deep valley (Fig. 1f-g and Supplementary Fig. 2). Similar to surface-immobilized MTs, DDB and yeast dynein were able to walk at the highest QD concentration tested on MT bridges (Fig. 1h). Interestingly, yeast dynein’s run length was 2.5-fold higher on MT bridges compared to surface-immobilized MTs at 12 QD μm^-1^ (p < 0.001, two-tailed t-test, Fig. 1e-h), suggesting that this motor bypasses obstacles more successfully by exploring the entire MT surface. However, we did not observe a significant improvement in kinesin motility on MT bridges in comparison to surface-immobilized MTs (Fig. 1e-h). Mobile fraction was reduced 75% at 1 QD μm^-1^ compared to the no obstacle condition and motility could not be detected at 7 QDs μm^-1^. These results suggest that kinesin is intrinsically limited by its ability to side-step to adjacent protofilaments when it encounters obstacles on an MT.

**Fig. 2.**
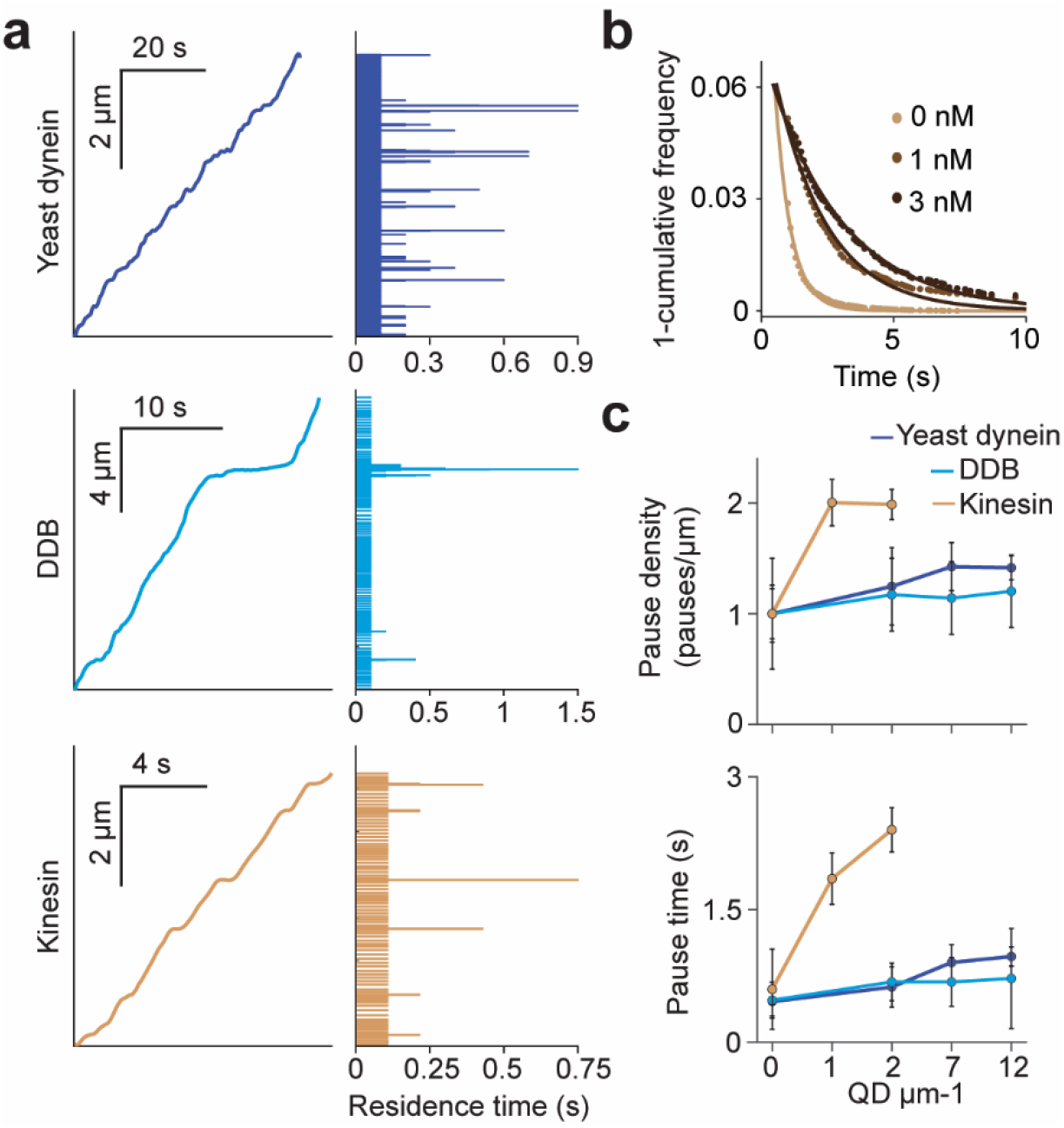
Kinesin motors pause longer and more frequently when encountering QD obstacles. **a** (Left) Representative traces of yeast dynein, DDB and kinesin in the absence of QD obstacles on surface-immobilized MTs. (Right) Residence times of the motors in each section of the traces. **b** Inverse cumulative distribution of kinesin residence times at different obstacle concentrations were fit to a single exponential decay. The residuals of that fit (shown here) are fit to a single exponential decay (solid line) to calculate the density and duration of kinesin pausing. c Density and durations of the pauses of the three motors. Pause densities are normalized to the 0 nM QD condition. Kinesin pausing behavior at 7 and 12 QDs μm^-1^ could not be determined, because the motor was nearly immobile under these conditions. From left to right, *n* = 535, 520, 158, 29 for yeast dynein, 511, 449, 391, 276 for DDB, and 570, 127, 112 for kinesin. Error bars represent SE calculated from single exponential fit to residence times.

### Kinesin motors pause when encountering an obstacle

We next investigated how obstacles affected the pausing behavior of motors. Even at the lowest QD density, most kinesin motors were immotile over the course of the recording, suggesting that kinesin has a high likelihood of permanently pausing when encountering an QD. Trajectories of the remaining processive motors were interspersed with frequent pauses (Supplementary fig. 3). We analyzed the trajectories of these motors before they permanently paused or dissociated from the MT and calculated the residence times of motors per distance traveled (Fig. 2a). Residence times were composed of two distinct states. A fast state corresponded to processive motility of the motor along MTs and a slow state represented transient pauses in motility (Fig. 2b). We calculated the density and length of pauses from the frequency and decay time of the slow state (Fig. 2b). Strikingly, kinesin pause density increased two-fold and pause time increased four-fold at 2 QDs μm^-1^. In contrast, pause density and duration of yeast and DDB were only modestly increased by the QD density (Fig. 2c). Transient pauses may correspond to detachment of kinesin when encountering an obstacle and reattachment to a nearby protofilament.^12^ However, this mechanism is not robust enough to efficiently bypass obstacles and kinesin motility stalls permanently usually after a few transient pauses in motility. Dynein also pauses frequently in the presence and absence of QD obstacles (Supplemental fig. 4). However, unlike kinesin, it rarely pauses permanently even at the highest density of QDs (Fig 2c).

### Multi-kinesin cargos can avoid obstacles

In cells, cargos are often carried by multiple motors, which increases collective force generation and enables transport of the cargo over longer distances^21–24^ as well as slightly higher velocities.^28^ We asked whether multiple motors can transport a cargo under conditions in which single motors are unable to walk along MTs. To test this, 500 nm cargo beads were coated with multiple kinesin or DDB motors (Fig. 3a). In the absence of QD obstacles, the beads were highly processive and did not detach until they reached the end of the MT. When the beads were incubated with a low concentration (50 nM) of kinesin motors, we detected processive motility of beads, albeit with frequent pausing, on MTs decorated with 7 QDs μm^-1^ (Fig. 3b). This was a density at which single kinesins were completely inhibited (Fig. 1e). However, motility of these beads was severely inhibited at 12 QDs μm^-1^. Surprisingly, when beads were incubated with a higher concentration (1.5 μM) of kinesin, their mobile fraction was unaffected by decoration of MTs with 12 QDs μm^-1^ (Fig. 3b-c, Supplementary fig. 5, Supplementary video 3). Similarly, multiple DDBs transported beads to the minus-ends of MTs regardless of the surface density of QDs with no decrease in mobile fraction (Fig. 3b-c). Both motors displayed a 25% decrease in velocity when exposed to 12 QDs μm^-1^ (p < 0.03, Fig. 3c). Therefore, while single kinesins are strongly affected by obstacles on MTs, team of kinesins can carry cargo beads over long distances as well as dyneins along MTs densely decorated with obstacles.

**Fig. 3.**
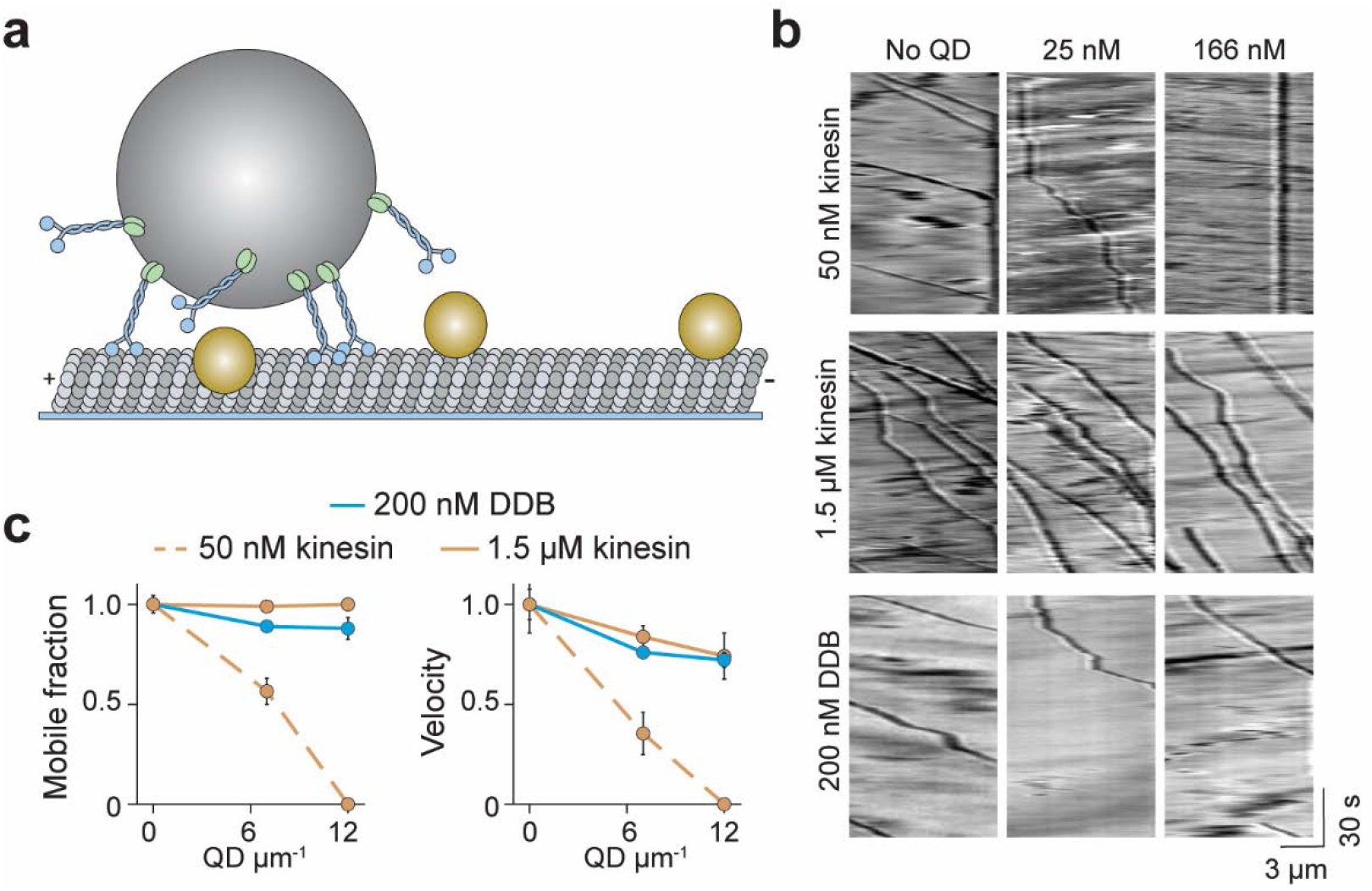
Cargos driven by multiple kinesins successfully bypass QD obstacles. **a** Schematic of bead motility driven by multiple motors along surface-immobilized MTs decorated with QD obstacles (not to scale). **b** Kymographs reveal the motility of beads coated with kinesin or DDB in the presence and absence of QD obstacles. **c** Mobile fraction and velocity of beads were normalized to the no QD condition (mean ± SD, three independent experiments). From left to right, *n* = 154, 189 bead traces for 50 nM kinesin, 323, 338, 336 for 1.5 μM kinesin, and 279, 184, 67 for DDB.

### Multi-motor cargos rotate around the MT to avoid large obstacles

Avoiding obstacles larger than a QD, such as a stationary organelle or intersecting cytoskeletal filament, may require a cargo to rotate to the other side of the MT before continuing forward movement. ^29^ *In vivo* studies have observed both anterograde and retrograde cargos to bypass stationary organelles^8^. It has been proposed that rotation of a cargo around the MT requires the presence of both kinesin and dynein motors on the cargo or the distortion of the lipid cargo.^26, 29, 30^ To test whether a single type of motor can rotate a rigid cargo around the MT, we tracked beads driven by multiple kinesins or dyneins on MT bridges. If the beads were positioned below the MT when they reached the end of the bridge, they were challenged to bypass the PDMS wall (Fig. 4a). Remarkably, we observed that nearly all of these beads rotated to the top of the MT with no forward motion before they continued along the MTs (77 ± 13% kinesin beads and 85 ± 5% dynein beads, mean ±SD, Fig. 4b-c, Supplementary Video 4 and 5). Of those events, 29% and 24% of kinesin- and dynein-driven beads, respectively, paused before moving forward (Fig. 4b-c), similar to intracellular cargos that encounter stationary organelles.^10^ This movement was clearly different from previously observed helical movement of kinesin- or dynein-driven cargos around the MT,^6, 31^ in which rotation is accompanied by forward translational movement. We observed no major differences in pausing or detachment behavior between kinesin- and dynein-driven beads. Surprisingly, multiple kinesins paused for a shorter period than dyneins when they encountered the wall (3.8 ± 0.2 s vs 7.8 ± 1.0 s, mean ± SEM, Fig. 4d). Therefore, bead rotation is unlikely to be driven by sidestepping of the motors to neighboring protofilaments. We concluded that cargos driven by multiple motors can bypass large obstacles by rotating around the circumference of the MT.

**Fig. 4.**
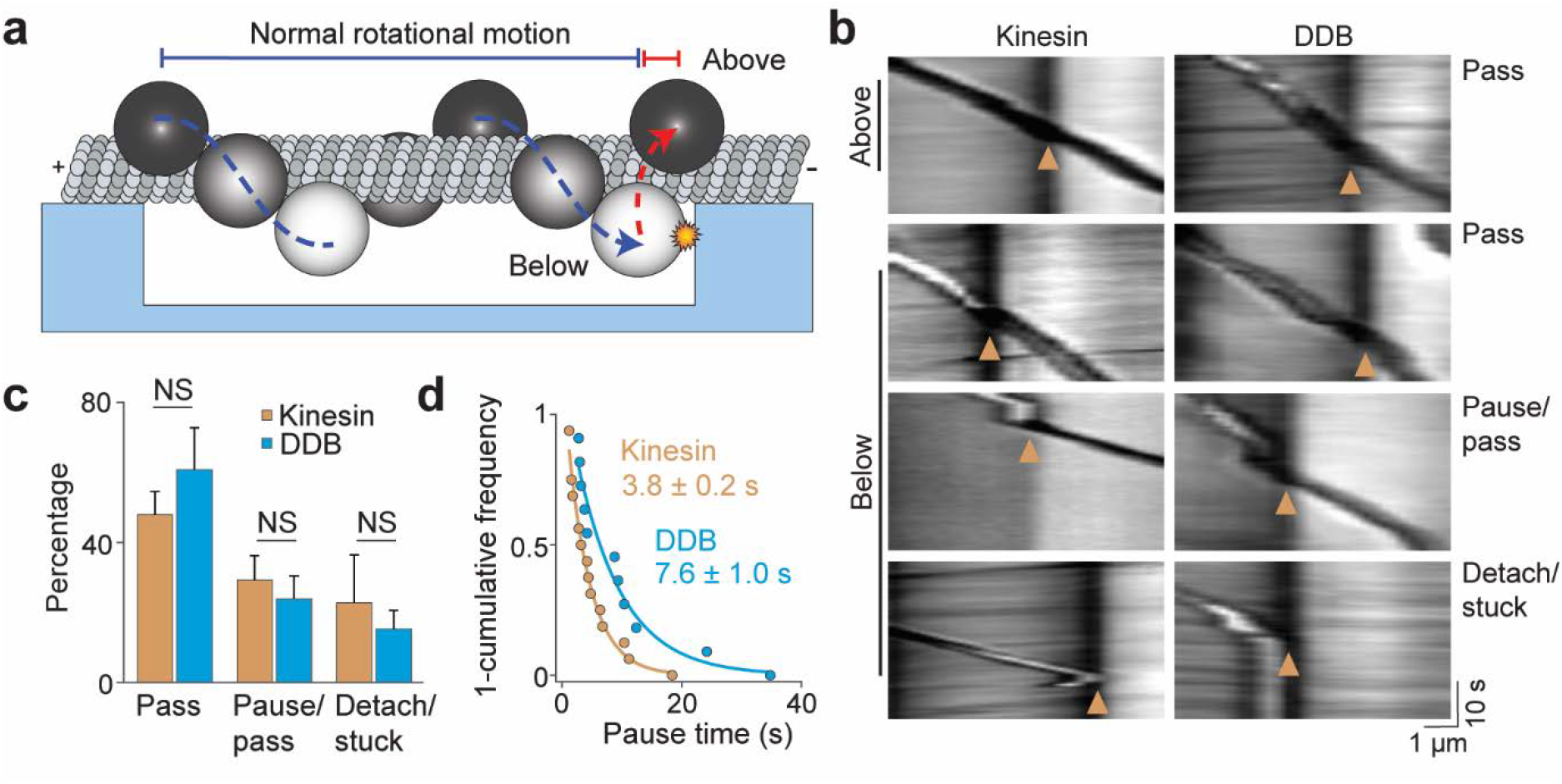
Cargos driven by multiple motors bypass large obstacles by rotating around the MT. **a** Schematic of multi-motor bead motility on MT bridges. The position of the bead in the *z* axis is determined from changes in bead intensity under brightfield illumination. If a bead is positioned below the MT when it reaches the PDMS wall, it must move to the top of the MT (red dotted curve) before continuing forward. **b** Kymographs reveal how beads driven by kinesins and DDBs move when they encounter the PDMS wall (Below). (Pass) The bead rotates around the MT as evidenced by light to dark transition in the bead intensity at the wall before continuing forward. (Pause/pass) The bead paused for more than 1 s at the wall before rotating around the MT and moving forward. (Detach/stuck) The bead failed to pass the wall and either detached or got stuck on an MT. **c** The percentage of pass, pause/pass and detach/stuck events for the beads positioned below the MT when they encounter the PDMS wall (mean ± SD, two independent experiments; NS: non-significant, two-tailed t-test; *n* = 122 for kinesin, and 81 for DDB beads). **d** Pause time distribution for kinesin and DDB beads. A fit to a single exponential decay (solid curves) revealed that pause duration of DDB-driven beads is longer than kinesin-driven beads (F-test, p < 0.001, *n* = 16 pauses for kinesin and 11 for DDB).

## Discussion

Despite the complexity of the cytoplasm, kinesin and dynein drive intracellular transport with remarkable efficiency towards MT ends. In this study, we investigated the ability of kinesin and dynein to bypass permanent obstacles using *in vitro* reconstitution and single-molecule imaging. These results show that motors utilize different mechanisms to bypass these obstacles. Despite their large size, single dyneins were highly capable of maneuvering around small obstacles on the MT. In contrast, single kinesins were strongly inhibited by such obstacles, as previously reported.^9, 12–14^ These results are likely a consequence of differences in the sidestepping ability of the two motors.^5, 6, 8^ Remarkably, multiple kinesins were able to bypass these obstacles as efficiently as dyneins. In the case of multi-motor cargos, we anticipate that kinesin motors are just as likely to get stuck at an obstacle as a single motor. However, the other motors driving the cargo exert a force on the stuck motor, causing its rapid detachment and the continued forward motion of the cargo.^32^

Multi-motor teamwork also proved beneficial for both types of motors when challenged by a large obstacle. Cargo beads driven by multiple kinesin or dynein motors were able to maneuver around a PDMS obstruction that blocked access to half of the MT surface. This behavior helps explain how cargos bypass large cellular roadblocks such as stationary organelles or intersecting cytoskeletal filaments. Recent measurements observed rotational motion in the trajectories of endosomal cargos carrying gold nanorods.^30^ In addition, correlative live-cell and super-resolution microscopy showed that rotational movement could be used by cargos to avoid steric obstacles.^29^ These studies proposed that rotational movement and off-axis stepping might result from having a mix of motors, such as kinesin-2 or dynein, on the cargo along with kinesin-1. While transient back-and-forth movement of a cargo may allow it to change the protofilament track, our results clearly show that tug-of-war between opposite polarity motors is not required to bypass large obstacles. Instead, cargo beads driven by multiple motors can switch to the other side of the MT surface by rotation, when only one type of a motor is active at a time. We also showed that fluidity of the cargo is not essential for this process.^29^ These results show that multiplicity of motors not only increases the collective force generation and the length of processive runs on an MT, but also enables motors to maneuver around obstacles in their path. Future studies are required to address how the motor copy number affects the ability of a cargo to bypass dynamic obstacles, such as MAPs.

## Acknowledgements

We are grateful to V. Belyy, A. Jack and Y. Ezber for helpful discussions, S.M. Luk for helium ion microscopy, P. Lum and N. Azgui at the Biomolecular Nanotechnology Center for help with nanofabrication, and A. Killilea for the mammalian cell culture.

## Author contributions

L.F. and A.Y. designed the study. L.F. purified the protein and performed all experiments. M.M.E. established the insect cell purification procedure. M.A.T. performed DDB experiments with obstacles. S.C. wrote the simulation program. L.F. and A.Y. wrote the manuscript.

## Supplementary Information

**Supplementary Fig. 1.**
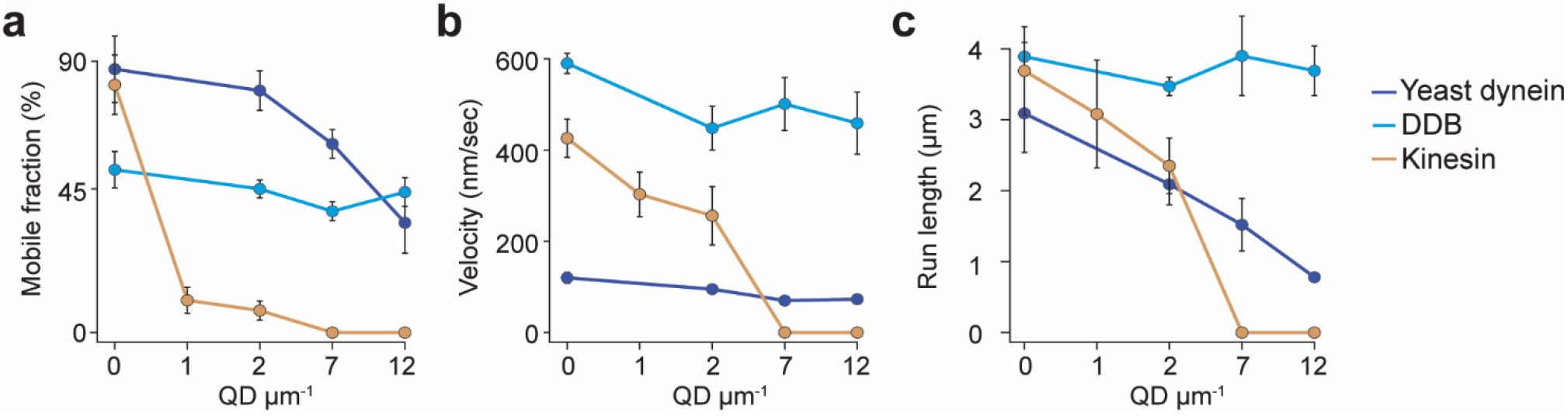
Analysis of single molecule trajectories on surface-immobilized MTs without normalization. **a** Mobile fraction, **b** velocity and **c** run length of single motors on surface-immobilized MTs in the presence of QD obstacles (mean ± SD, three independent experiments). From left to right, *n* = 271, 423, 405 for kinesin, 315, 407, 197, 168 for yeast dynein, and 636, 502, 356, 509 for DDB.

**Supplementary Fig. 2.**
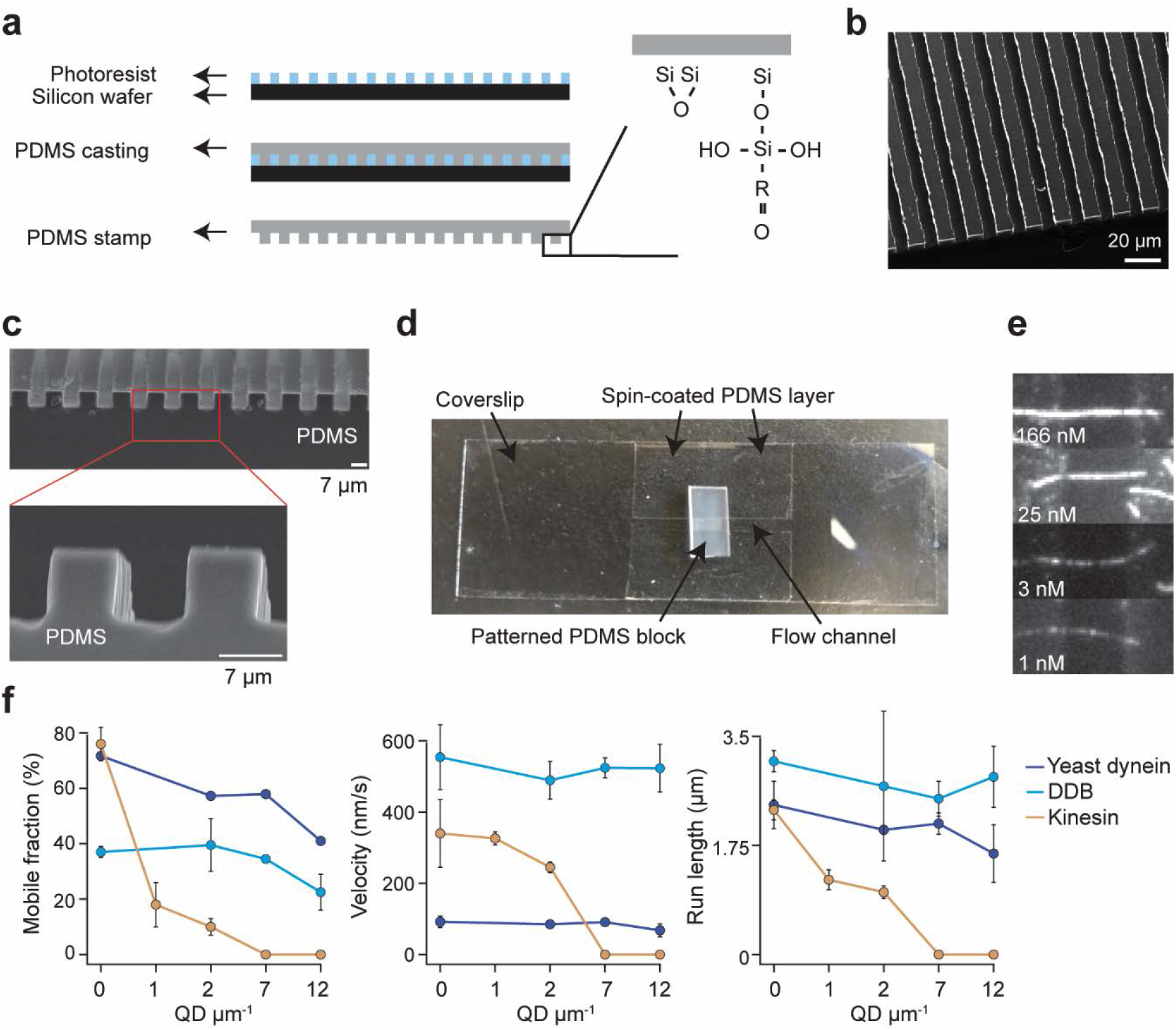
Chamber design and raw data of single molecule motility along MT bridges. **a** Workflow for bridge microfabrication. Photoresist is spun and patterned on silicon wafer. PDMS is then cast on top of the photoresist and silanized to produce a reactive surface **b** Helium ion microscopy shows top view of patterned PDMS. **c** Scanning electron microscopy shows the side view of patterned PDMS (top) and the zoomed view of this image reveals that the walls have sharp edges (bottom). d Image of the flow chamber used for experiments. **e** Example fluorescent images of QD585 obstacles on MT bridges at different QD concentrations. **f** Mobile fraction, velocity and run length of single motors on MT bridges without normalization (mean ± SD, two independent experiments). From left to right, *n* = 199, 187, 106 for kinesin, 129, 107, 163, 135 for yeast dynein, and 192, 206, 330, 276 for DDB.

**Supplementary Fig. 3.**
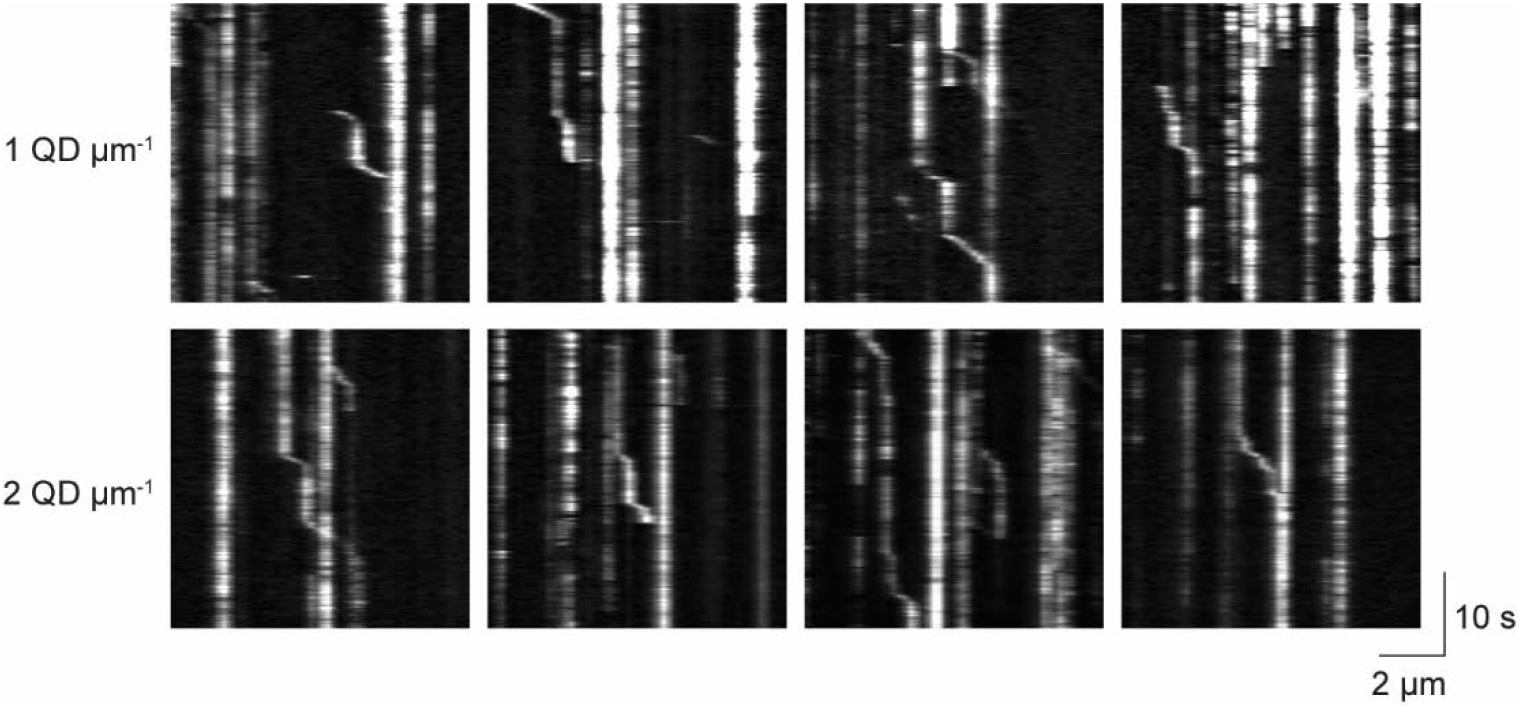
Kinesin pauses in the presence of QD obstacles. Representative kymographs reveal frequent pauses in kinesin motility in the presence of 1 QD μm^-1^ (top row) or 2 QD μm^-1^ (bottom row). Most pauses were permanent over the course of recording. Processive traces interspersed with transient pauses were used in pause analysis in Figure 2.

**Supplementary Fig. 4.**
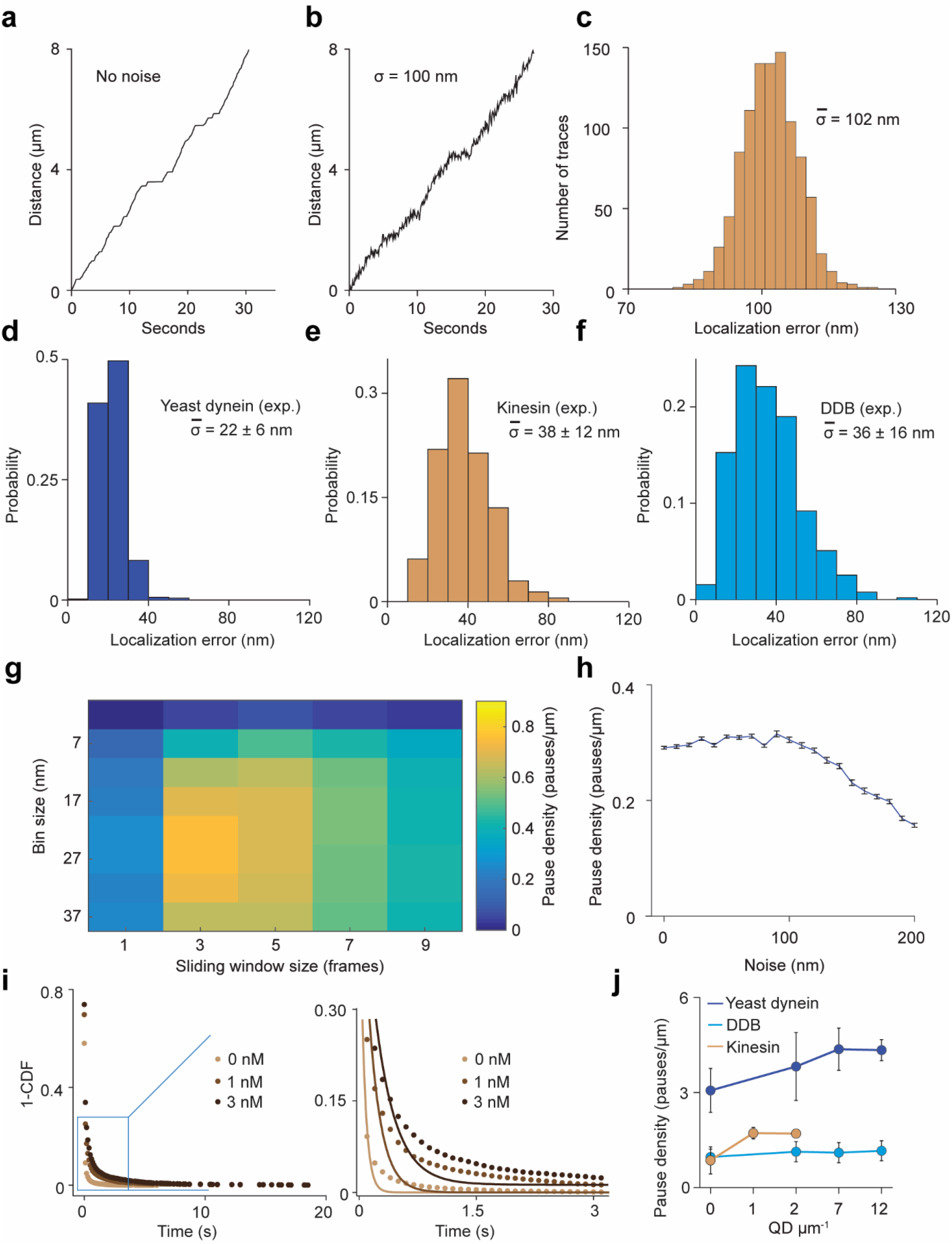
Simulations for the pause analysis. **a** An example trajectory simulated with a pause density of 0.8 μm^-1^ in the absence of tracking noise (see Methods for the parameters used to generate these trajectories). **b** An example trajectory simulated with 100 nm root mean squared (RMS) noise. (C) Localization error calculated for 1,000 simulated traces closely agrees with 100 nm noise added to the traces. **d-f** Localization error calculated for 500 experimental traces of the three motors. **g** Optimization of the bin size and sliding window size for the pause analysis. Noisy traces were simulated using 0.8 μm^-1^ pause density, downsampled with a given window size (the number of data points) and residence time was calculated for a given bin size (distance traveled by motor). The analysis revealed that pause density was slightly underestimated even under optimum conditions. The combination of bin size and window size that resulted in the highest pause density was used to analyze experimental traces. **h** Traces were simulated with a pause density of 0.3 μm^-1^. Calculated pause density from simulations was insensitive to the 0-100 nm added tracking noise. The density of detected pauses decreases at higher noise. **i** Pause density and duration were determined from residence time histograms through a two-step process. (Left) All non-zero residence times were fit to a single exponential distribution. (Right) Zoomed view of the blue rectangle on the left. The residuals of this fit (plotted in Fig. 2B) were fit to a single exponential decay to determine pause time and density. **j** The pause density analysis of single motors on surface-immobilized MTs without normalization. From left to right, *n* = 535, 520, 158, 29 for yeast dynein, 511, 449, 391, 276 for DDB, and 570, 127, 112 for kinesin. Error bars represent SE calculated from single exponential fit to residence time histograms.

**Supplementary Fig. 5.**
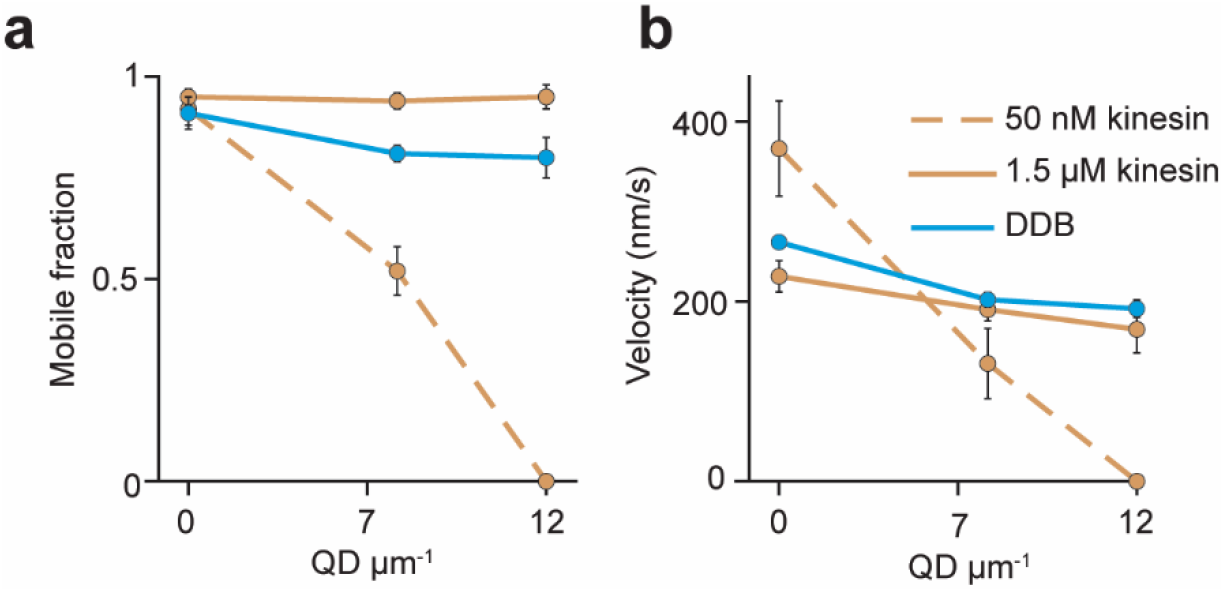
The analysis of beads driven by multiple motors on surface-immobilized MTs without normalization. **a** Mobile fraction and b velocity of beads driven by multiple motors on surface-immobilized MTs in the presence of QD obstacles (mean ± SD, three independent experiments). From left to right, *n* = 154, 189 traces for 50 nM kinesin, 323, 338, 336 traces for 1.5 μM kinesin, and 279, 184, 67 traces for 200 nM DDB.

## Methods

### Protein expression and purification

A human kinesin-1 coding sequence expressing amino acids 1-560 was fused to GFP and HaloTag on the C-terminus (hK560::GFP::HaloTag).^33^ The N-terminus of yeast cytoplasmic dynein was replaced with a HaloTag and a GST dimerization tag (HaloTag-GST-Dyn1_331kDa_, consisted of amino acids 1219-4093 of the dynein heavy chain).^34^ A full length human cytoplasmic dynein construct consisted of the dynein heavy chain tagged with an N-terminal SnapTag, cloned into a pOmniBac vector, and fused to a plasmid that containing dynein intermediate chain, light intermediate chain and three different light chains (Tctex, Roadblock and LC8), as described^18^. The BicD2N construct consisted of GFP fused to the first 400 amino acids of mouse Bicd2^18^.

### Kinesin purification

Rosetta cells transformed with kinesin plasmid were grown in a 5 mL culture overnight. This culture was added to 1 L of LB media and grown for 3 hours until the OD600 reached 0.7. Cells were induced with fresh 100 μM IPTG, put on ice until the temperature reached 20 degrees and incubated overnight at 20 degrees at 180 RPM. After harvesting cells at 4785 RCF for 15 minutes in a JLA 8.1 rotor, 500 mL cell pellets were incubated with 40 mL lysis buffer (50 mM sodium phosphate monobasic pH 8.0, 250 mM sodium chloride, 2 mM magnesium chloride, 20 mM imidazole, 1 mM ATP, 10 mM betamercaptoethanol (BME), 1 mM phenylmethylsulfonyl fluoride (PMSF). Cells were lysed with a sonicator and spun in a Ti70 rotor at 117734 RCF for 30 min. Supernatant was incubated with 6 mL of washed Ni-NTA bead slurry (HisPur, Thermo Scientific) for 1 hour at 4 degrees. Beads were collected in a BioRad column and washed in wash buffer (50 mM sodium phosphate monobasic pH 6, 250 mM sodium chloride, 1 mM magnesium chloride, 20 mM imidazole, 100 μM ATP, 10 mM BME). Protein was eluted in elution buffer (50 mM sodium phosphate monobasic pH 7.2, 250 mM sodium chloride, 1 mM magnesium chloride, 500 mM imidazole, 100 μM ATP, 10 mM BME) and snap frozen in 20% glycerol.

### Yeast dynein purification

Yeast cells were cultured on YPAD plates for 2-3 days. A 10 mL culture was grown overnight in YP media with 1 mL of 25% dextrose + 0.04% adenine supplements at 30 degrees overnight. 2 mL of the culture was then added to 100 mL of 1.25x YP media supplemented with 10 mL of 20% raffinose. After 9 hours of growth, the entire culture was added to 1.8 L of YP media supplemented with 200 mL of 20% (w/v) galactose. Cells were cultured at 30 degrees with shaking (200 rpm) overnight until the OD600 reached 1.5. After harvesting cells at 4785 RCF for 15 minutes in a JLA 8.1 rotor, cells were frozen dropwise and lysed while frozen in a coffee grinder. 50 mL of lysis buffer (30 mM HEPES pH 7.4, 50 mM potassium acetate, 2 mM magnesium acetate, 1 mM EGTA, 10% glycerol, 1 mM dithiothreitol, 100 μM ATP, 1 mM PMSF) was added to a 1 L yeast pellet.

Cells were spun at 360562 RCF for 45 min in a Ti70 rotor. Supernatant was incubated with washed IgG beads (IgG Sepharose 6 Fast Flow, GE Healthcare) for 1 hour with gentle rolling. Beads were collected using a BioRad disposable column, washed with wash buffer (lysis buffer with 125 mM KCl) and TEV buffer (10 mM Tris pH 8, 150 mM KCl, 10% glycerol, 1 mM tris(2-carboxyethyl)phosphine, 100 μM ATP, 1 mM PMSF). Beads were transferred to an Eppendorf tube and eluted with TEV protease for 1 hour. Beads were then spun down and the supernatant was snap frozen in 20% glycerol.

### Human dynein, human BicD and porcine dynactin purification

Human dynein, BicD2N and dynactin were purified as previously described.^35^ Further information can also be found on Invitrogen's Bac-to-Bac Baculovirus Expression System Guide (Invitrogen). Briefly, plasmids containing genes of interest were transformed into DH10Bac competent cells and plated on LB agar plates with kanamycin, gentamycin, tetracycline, Blue-gal and isopropyl beta-D-1-thiogalactopyranoside. An overnight culture of a colony grown in 2X YT media with kanamycin, gentamycin and tetracycline and the bacmid was purified from these cells. Cells were lysed and neutralized using Qiagen miniprep buffers P1, P2 and P3. DNA was then precipitated with isopropanol and spun down for 10 min at 13,000 RCF at 4 °C. The DNA pellet was washed three times with 70% ethanol, air dried and resuspended in Qiagen's EB buffer.

Bacmid was used within a few days for transfecting Sf9 cells. All insect cell culture was courtesy of Berkeley's Cell Culture Facility. 2 mL of Sf9 cells at 500,000 cells/mL was aliquoted into a 6-well dish and allowed to attach for 10 minutes. 1 microgram of bacmid DNA was diluted in ESF 921 media (Expression systems, no antibiotic or serum), mixed with 6 μL of Fugene HD transfection reagent (Promega) and incubated for 15 minutes at room temperature. Media on the cells was removed and replaced with 0.8 mL of ESF 921 media. The Fugene/DNA mix was added dropwise on the cells. The dish was sealed with Parafilm and incubated for 72 h. 24 h into this incubation, 1 mL of extra ESF 921 media was added to the cells. After removing the media and spinning, 1 mL of the supernatant (P1 virus) was added to 50 mL of Sf9 cells at a density of 1 million cells/mL. Following a 72 h incubation, the media was spun down and the supernatant (P2 virus) was harvested. 10 mL of the P2 virus was used to infect 1 L of Sf9 cells at 1 million cells/mL and expression proceeded for 72 hours. Cells expressing protein of interest were harvested at 4000 RCF for 10 min and resuspended in 50 mL lysis buffer (50 mM HEPES pH 7.4, 100 mM NaCl, 10% glycerol, 1 mM DTT, 100 μM ATP, 2 mM PMSF and 1 tablet of protease inhibitor cocktail). Lysis was performed using 15 loose and 15 tight plunges of a Wheaton glass dounce. Lysate was clarified using a 45 minute, 360562 RCF spin in a Ti70 rotor and incubated with 2 mL IgG beads (IgG Sepharose 6 Fast Flow, GE Healthcare) for 2 h. Beads were washed with lysis buffer and TEV buffer (50 mM Tris pH 7.4, 150 mM potassium acetate, 2 mM magnesium acetate, 1 mM EGTA, 10% glycerol, 1 mM dithiothreitol, 100 μM ATP). Beads were then collected and incubated with TEV protease overnight to elute the protein. Finally, protein was concentrated and frozen in a 20% glycerol solution.

### Glass silanization

Glass slides were functionalized with aminopropyltriethoxysilane (APTES) and glutaraldehyde to allow for covalent attachment of MTs, as described previously.^36^ APTES (Sigma, 440140) aliquots were prepared in 5 mL cryo tubes (Corning, 430656) using glass pipettes, capped under nitrogen atmosphere, flash frozen upright in liquid nitrogen and stored in −80 °C. Glass slides were sonicated in a 2% Mucasol (Sigma, Z637181) prepared in hot water and then rinsed thoroughly in water. Slides were then baked on a hot plate (Benchmark, BSH1002) to remove excess water for 5 min. To create functional silanol groups, slides were treated with oxygen plasma (PETS Reactive Ion Etcher) at 200 mTorr oxygen, 55 W for 1 min. Slides were rinsed briefly in acetone (Sigma, 270725) and immersed in a 2% (v/v) APTES in acetone for 1 min. APTES aliquots were added to acetone before warming to room temperature. After silane treatment, the slides were rinsed in acetone, and baked on a 110 °C hot plate for 30 min. To remove silane unbound to the glass, slides were sonicated sequentially in ethanol (Sigma, 459828) and water for 5 min. Following this treatment, slides were again baked at 110 °C for 30 min. An 8% glutaraldehyde solution (Fisher Chemical, G1511) was prepared in water and 1 mL drops of the solution were made on Parafilm. Slides were incubated functional-side down on the glutaraldehyde solution drops for 30 min in a sealed container. Finally, the slides were washed and sonicated in water for 10 s to remove loosely absorbed glutaraldehyde and stored in a sealed container at room temperature up to 1 week.

### Labeling

Motors were labelled with QDs modified with a HaloTag or SnapTag ligand. Amino-PEG-QDs (Thermo, Q21521MP) were labelled with HaloTag ligand by reaction with N-hydroxysulfosuccinimide reactive Halo-Tag ligand (Promega, P6751) or Snap-Tag ligand for 30 min at room temperature. QDs were then exchanged into 25 mM borate pH 8.5 using Amicon 100K centrifugal filters and stored in that buffer. 2 μM of these QDs were mixed with 100-500 nM motors fused with a SNAPTag or HaloTag for 10 min on ice.

### Motility assays

Tubulin was purified and labelled with biotin or fluorophores as described.^36, 37^ The final percentage of biotin on the MT was less than 5%. To perform motility assays, biotinylated MTs were diluted in BRB80 (80 mM PIPES pH 6.8, 1 mM EGTA, 1 mM magnesium chloride) supplemented with 10 μM taxol and flowed into a chamber made with two pieces of double-sided tape between an APTES-silanized slide and an unmodified coverslip. Chamber was then passivated with BRB80 supplemented with 1 mg/mL casein (Sigma, C5890), 1 mM DTT and 10 μM taxol. The chamber was incubated with different dilutions of streptavidin-coated QDs (Invitrogen, Q10111MP). BRB80 (above), DLB (30 mM HEPES pH 7.2, 2 mM magnesium chloride, 1 mM EDTA, 10% glycerol) and MB (30 mM HEPES pH 7.0, 5 mM MgSO_4_, 1 mM EGTA) buffers were used for assaying the motility of kinesin, yeast dynein and DDB, respectively. Motor-QD mixtures were flowed into the chamber, bound to the MT and washed to remove unbound motor and QD. Finally, motor-specific buffer supplemented with 1 mg/mL casein, 1 mM tris(2-carboxyethyl)phosphine (TCEP), 100 μM ATP, glucose oxidase, catalase and 0.8% dextrose was flowed into the chamber. Run length and velocity were determined by selecting the beginning and end of each trace in ImageJ. Cumulative frequencies of run lengths were fit to a single exponential function. Mobile fraction was calculated by dividing the number of moving motors over the total number of motors observed in the kymograph.

### Microscopy

Microscopy was performed using a custom-built fluorescence microscope equipped with a Nikon Ti-E Eclipse microscope body, a 40X 1.15 NA long-working-distance water immersion objective (Nikon, N40XLWD-NIR), and a perfect focusing system.^8^ The sample position was controlled by using automated microscope stage (Microstage 20E, MadCityLabs). The sample was excited in the epifluorescence mode using 488, 561 and 633 nm laser beams (Coherent). Fluorescence image was split onto two channels using 0ptoSplit2 (Cairn instruments) and detected by an electron-multiplied CCD Camera (Andor Ixon, 512×512 pixels).

### Single-molecule tracking and pause analysis

Single particle tracking was performed using Utrack.^38^ Tracks were split into 1D motion along the long axis of the MT and perpendicular direction as described.^8^ All tracks were manually reviewed to exclude tracks with jumps greater than 100 nm. Localization error was calculated using high pass filtering of the trajectories by calculating 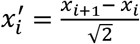, where *x* is the position of the probe along the MT axis and *i* is the frame number. This operation omits unidirectional motility and pauses at lower frequencies, leaving only the Gaussian noise associated with the trace. *σ_x′_* is defined as the localization error. Under our imaging conditions, localization error of QDs was typically between 20-40 nm (Supplementary Fig. 3). Pause analysis was performed as described.^39^ Briefly, the track was divided into distance bins and the residence time within each bin was calculated. All non-zero residence times were fit to a single exponential decay. The residuals were then fit to a single exponential decay. The decay time and amplitude of this second fit were defined as the average pause time and pause density, respectively.

### Simulations

Optimum running window averaging and bin size were calculated from simulated traces generated in MatLab. Experimentally determined noise, velocity and pause distributions were used to generate traces (Supplementary Fig. 3). A particle takes 8 nm unidirectional steps with exponentially distributed dwell times of 0. 015 and 0.59 s, mimicking the characteristics times of processive motility and pausing of kinesin motors, respectively. The trace was then resampled to the imaging rate of 10 Hz. Random Gaussian noise was added to each position to introduce localization error. Simulations were used to determine whether the localization error in traces interferes with pause detection (Supplementary fig. 3h).

### Microfabrication

PDMS bridges were generated using soft lithography.^40, 41^ Briefly, SU-8 2010 negative photoresist (Microchem) was spun on to a silicon wafer to 10 μm thickness. After a soft bake, photoresist was exposed to UV light through a patterned, film photomask (Fine Line Imaging) on an OAI 200 Lithographic Mask Aligner. The pattern was developed after a hard bake using SU-8 developer (Microchem). To render the surface less adhesive, the master was treated with trichloro(1H,1H,2H,2H-perfluorooctyl)silane vapor. Sylgard 184 (Dow Corning) base and curing agent were mixed in a 10:1 ratio by mass and degassed. After pouring over the master, the PDMS was cured for 1 h at 80 °C (Supplementary Fig. 2). Features were confirmed with helium ion microscopy and scanning electron microscopy. The PDMS was then removed from the mask and baked for an additional 1 week at 80 °C. To extract low molecular weight species that diffuse to the surface and alter surface chemistry,^42^ PDMS was incubated, in sequence, with triethylamine, ethyl acetate, and acetone and allowed to dry overnight in an oven.

### MT bridges

Silanization of PDMS was similar to glass with a few modifications. Ethanol was used as the solvent rather than acetone, as it is less likely to swell the PDMS.^43^ The patterned surface was plasma oxidized at 50 W and 200 mTorr for 1 min. The slab was then immediately immersed in a 5% (v/v) solution of APTES in HPLC-grade ethanol (Sigma, 459828) for 20 min. After rinsing in 95% ethanol/water, the PDMS was baked 40 min on a hot plate. To remove unbound silane, PDMS was rinsed in pure ethanol and then water, and baked for an additional 40 min. Finally, PDMS was incubated for 1 h in an 8% glutaraldehyde solution. Excess glutaraldehyde was removed by rinsing in water and the functionalized PDMS was stored at room temperature for 1 week. To create a flow chamber, uncured PDMS was spin-coated on a coverslip to a thickness of 100 μm. After baking, a channel was cut in the PDMS-coated coverslip and the surface was plasma oxidized. The PDMS block with the bridge pattern was placed functional-side down on the coverslip, creating a flow cell (Supplementary fig. 2). Motility assays were performed as described for single-molecule assays and the sample was imaged using an 1.4 NA oil-immersion condenser (Nikon) under brightfield illumination.

### Anti-GFP coating beads

Latex beads were coated with anti-GFP antibody as described.^33^ To prevent clumping of the beads when incubated with high concentrations of motors, we used “CML” beads (ThermoFisher, C37481), which have high density of carboxyl groups that facilitates charge repulsion. 200 μL of 0.5 μm diameter CML beads (4% solids) were washed three times in activation buffer (10 mM MES, 100 mM sodium chloride, pH 6.0) by centrifugation for 6 min at 7,000 g. Final resuspension was in 200 μL activation buffer. Beads were then sonicated for 1 min in a bath sonicator (Vevor). Separately, fresh 4 mg/mL solutions of EDC (1-ethyl-3-(3-dimethylaminopropyl)carbodiimide hydrochloride, ThermoFisher) and Sulfo-NHS (N-hydroxysulfosuccinimide) were prepared in water. 5 μL each of fresh EDC and Sulfo-NHS solutions were added to the beads. Beads were sonicated for 1 min and nutated for 30 min at room temperature. After three washes in PBS, beads were resuspended in 200 μL PBS and mixed with 200 μL 0.4 mg/mL anti-GFP antibody overnight with nutation. The beads were passivated by incubating with 10 mg/mL bovine serum albumen (BSA) overnight. Finally, beads were washed 5 times in PBS and stored at 4 °C with 1 mg/mL BSA supplement.

### Bead motility assay

For bead motility assays, anti-GFP beads were diluted two-fold in water and sonicated for 1 min to disperse the beads. GFP-tagged kinesin motors were incubated with beads for 10 min. Excess motor was then washed from the beads by diluting the mixture into 100 μL BRB80 supplemented with 1 mg/ml casein (BRB-C) and centrifugation at 8,000 *g* for 3 min. The supernatant was removed and the pellet was resuspended in 15 μL BRB-C supplemented with 1 mM TCEP, 100 μM ATP, glucose oxidase, catalase and 0.4% dextrose. Beads were then flowed into a flow chamber after surface-immobilization of biotinylated MTs. DDB experiments were performed in a similar fashion with a few exceptions. The GFP handle was on the cargo adaptor (BicD2N-GFP). We used a dynein mutant that does not form the autoinhibited phi-conformation^35^ to facilitate assembly of the DDB complex. 1 μL each of 1 μM human dynein complex, pig brain dynactin and BicD2N-GFP were mixed at a 1:1:1 molar ratio and incubated for 15 min before mixing with the beads. The mixture was pelleted at 8,000 g, resuspended in 15 μL MB supplemented with 1 mg/mL casein, 1 mM TCEP, 100 μM ATP, glucose oxidase, catalase and dextrose and added to the flow chamber.

### Statistics and reproducibility

Each measurement was performed with at least three independent replicates, and the exact number of repetitions is reported for each experiment. Each statistical analysis method is explicitly stated in the main text and/or figure legend.

### Data availability

All data that support the conclusions are available from the authors on request.

### Supplementary Video legends

**Supplementary Video 1: Motility of DDB motors is not strongly affected by the QD obstacles**. DDB motors were labeled with QD655 at their N-termini. Single-molecule motility of DDB in the presence of no obstacles (top) or 25 nM QD585 obstacles (bottom) on surface-immobilized MTs. The fluorescence signal of QD585 obstacles was collected in a separate channel, and not displayed. The sample was excited with 1.7 kW cm^-2^ 488 nm laser beam under the epifluorescence mode. Images were acquired at 10 Hz.

**Supplementary Video 2: Kinesin motors walk processively in the absence of QD obstacles, but motility was completely impaired in the presence of 25 nM QDs**. Kinesin motors were labeled with QD655 at their C-termini. Single-molecule motility of kinesin in the presence of no obstacles (top) or 25 nM QD585 obstacles (bottom) on surface-immobilized MTs. The fluorescence signal of QD585 obstacles was collected in a separate channel, and not displayed. The sample was excited with 1.7 kW cm^-2^ 488 nm laser beam under the epifluorescence mode. Images were acquired at 10 Hz.

**Supplementary Video 3: Beads driven by multiple kinesins move processively at high obstacle concentrations**. 500 nm diameter beads were labeled with (1.5 μM) kinesin. Beads move along surface-immobilized MTs in the presence of no obstacles (left) or 166 nM obstacles (right). Boxes highlight the processive motility of beads along a surface-immobilized MTs (not labeled). In addition, freely diffusing beads in the chamber come in and out of focus during imaging. Images were acquired at 10 Hz under brightfield illumination.

**Supplementary Video 4: Beads driven by multiple kinesins bypass the PDMS wall**. The box highlights the processive motility of a bead driven by multiple kinesins on a MT bridge (unlabeled) suspended over the PDMS ridges. The valley (dark) is in the center of the movie while the PDMS ridges (light) are on either side. As the bead reaches the PDMS wall (light/dark interface), the bead intensity shifts from light to dark before the bead continues to walk on the PDMS ridge. Images were acquired at 10 Hz under brightfield illumination.

**Supplementary Video 5: Beads driven by multiple DDBs bypass the PDMS wall**. The box highlights the processive motility of a bead driven by multiple DDBs on a MT bridge (unlabeled) suspended over the PDMS ridges. The valley (dark) is in the center of the movie while the PDMS ridges (light) are on either side. As the bead reaches the PDMS wall (light/dark interface), the bead intensity shifts from light to dark before the bead continues to walk on the PDMS ridge. Images were acquired at 10 Hz under brightfield illumination.

